# Diversity of Novel Bacteriophages Infecting Ammonia-Oxidizing Bacteria

**DOI:** 10.64898/2026.05.02.722434

**Authors:** Aaron A.B. Turner, Miranda Stahn, Andrew Millard, Dominic Sauvageau, Lisa Y. Stein

## Abstract

Agriculture is a major source of anthropogenic greenhouse-gas emissions and the largest source of nitrous oxide (N2O), an extremely potent greenhouse gas and ozone-depleting agent. Soil N_2_O emissions are largely driven by nitrification, in which ammonia-oxidizing microorganisms catalyze the oxidation of ammonia to nitrite. Nitrification not only mediates N_2_O fluxes but also reduces fertilization efficiency and contributes to eutrophication through nitrate leaching.\ Bacteriophage (phage)-based control of microbial communities is gathering interest; however, phages infecting ammonia-oxidizers are mostly uncharacterized, with only one lytic phage having been described, limiting the potential for developing phage-mediated nitrification control. Here, we characterized the largest set of cultivated phages infecting ammonia-oxidizing bacteria (AOB) to date: 45 dsDNA phages collected from urban wastewater, infecting four AOB species, with 16 demonstrating cross-genus host ranges and capable of eliminating nitrification activity in liquid cultures. Phylogenetic and taxonomic analyses revealed six proposed families of *Caudoviricetes* and several monophyletic clades, likely representing higher-level lineages. Structure-guided genome annotation revealed a diverse collection of auxiliary metabolic genes encoded by phage, ranging from a complete ABC transporter cassette to a large antimicrobial resistance gene cluster. These results unveil previously unrecognized diversity of cultivated AOB phages and their potential to alter host physiology. Our data describes a broad taxonomic and functional repertoire of cultured AOB phages, suggesting that viruses play a significant and complex role in nitrification. Moreover, we outline an effective methodological framework for isolating AOB phages from environmental samples. These results reframe our understanding of environmental nitrification and enable intensified cultivation, characterization and use of phages for its control.

## MAIN TEXT

Climate change is an existential threat to Earth’s ecosystems, with irreversible effects already in force^1^. Without greenhouse gas (GHG) emissions mitigation, temperatures will continue to rise, and climate effects are expected to worsen. Agriculture comprises ca. 40% of land space while also being the foremost source of nitrous oxide (N_2_O) emissions according to the World Bank data on agricultural land^2^. N_2_O is a potent GHG with a heat-carrying capacity of ∼273 CO_2_-eq over 100 yr, while also being the dominating ozone-depleting agent^3,4^. The majority of N_2_O arises due to the overuse of nitrogen fertilizers (ammonia and urea-based), which commonly exceed crop demands, causing an imbalance in the nitrogen cycle^1,5^.

Soil N_2_O emissions largely result from microbial activities through the oxidation of ammonia to nitrate via nitrification and the reduction of nitrate to nitric oxide (NO), N_2_O, and N_2_ via denitrification^6,7^. The rate-limiting step of the N-cycle is the oxidation of ammonia to nitrite by ammonia-oxidizing microorganisms (AOM), with ammonia-oxidizing bacteria (AOB) as direct producers of N_2_O through nitrifier-denitrification activity and abiotic reactions of their metabolic intermediates^6,8^. Nitrate production, facilitated mainly by nitrite-oxidizing bacteria (NOB), drives eutrophication from its leaching into water systems^9^. Eutrophication causes substantial ecological damage and, in fact, is one of the most significant water quality problems globally. Moreover, nitrification is a major cause of low nitrogen use efficiency by crops, causing 50-70% losses in applied nitrogen^10,11^.

Developing an effective means of AOB control while bolstering fertilization and nitrogen use efficiency of crops is paramount in decreasing the climate and ecological impacts of agriculture. There are emerging efforts to achieve this, one of which is to leverage bacteriophages (phages) against soil AOB communities. However, knowledge of AOB phages is extremely limited, with only one strictly virulent phage – *Nitrosomonas* phage ΦNF-1 – having been isolated to date^12^. Although phages have the potential to control AOB populations including phage ΦNF-1^13^, diverse and extensive collections of phages are required to develop effective strategies; for example for the development of multi-phage cocktails which have significant advantages over single-phage treatments in other applications^14^. Moreover, the diversity and functional characteristics of AOB phages remain underexplored, particularly regarding their potential influence on shaping host physiology, community dynamics and the nitrification process.

## RESULTS

### Isolation of phages infecting AOB

As AOB, and by extension their cognate phages, have low relative abundance in wastewater influent compared to other bacteria and phages^15^, a filtration-based screening followed by liquid culture enrichment approach modified was implemented^16^ (Fig. 1). This approach enables screening of low-abundance phages – preliminary experiments for the filtration-based screening performed with phage T4 and *Escherichia coli* showed consistent recovery of as few as one phage virion per 250 mL screening volume (Fig. S1) –, reuse of raw sample against multiple pages, and isolation on hosts that do not grow on solid media.

**Figure 1.**
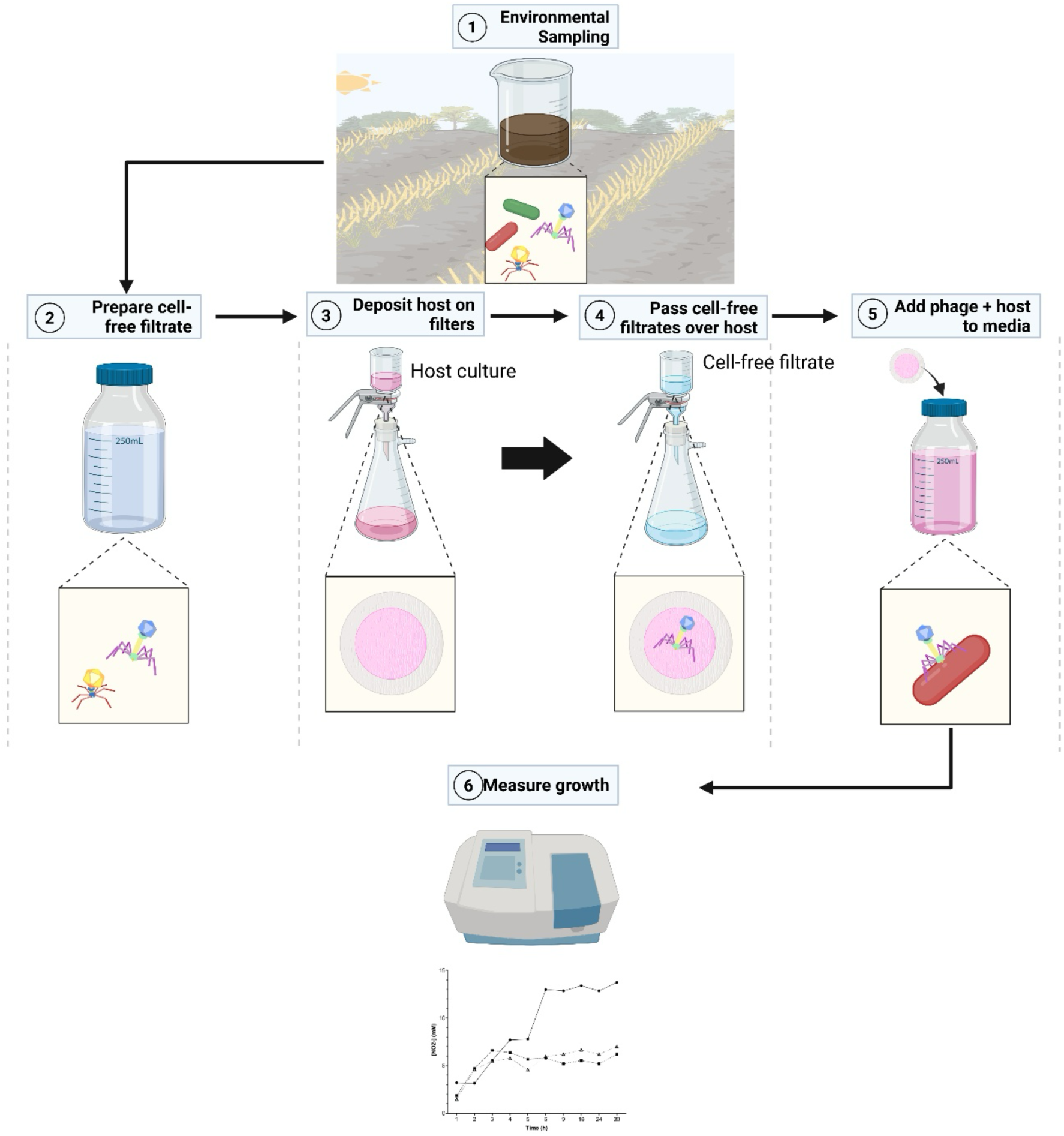
Modified filtration method for low-abundance phage screening. Environmental samples are filtered to generate cell-free filtrate samples. To screen for phages, host cells are deposited on filters (cell-deposited filters), and cell-free filtrates are passed over to capture phages capable of binding to the host. Filters are then transferred to AOB medium, incubated and cellular activity is measured by nitrite production. Successful infection was indicated by early cessation of nitrite production and decreased nitrite yield compared to control cultures undergoing the same experimental steps; these are filtered and designated as cell lysates.

Phages infecting hosts from the two major genera of AOB, *Nitrosospira* and *Nitrosomonas*, were identified by sequentially passing raw influent from two municipal wastewater treatment plants against cells of *Nitrosospira multiformis* ATCC 25196, *Nitrosospira briensis* C-128, *Nitrosomonas europaea* ATCC 19718, and *Nitrosomonas communis* Nm2 previously deposited on filters. Infection was assessed through the early cessation of nitrite production, compared to control cultures, when treated filters were added to fresh AOB medium.

12 samples demonstrating early cessation of nitrite production were harvested and designated as cell lysates for phage characterization. Transmission emission microscopy (TEM) micrographs confirmed the presence of phage particles in each of these 12 lysates, with most having mixed phage populations, while phages ripa and ripi1-4 representing the only pure isolate following initial screening, with rov1 and roa1 later isolated by host range (Fig. S2; Table S1). Phage morphology encompassed the three major morphotypes of *Caudoviricetes* phages. Notably, *N. europaea* ATCC 19718 and *N. briensis* C-128 phages were restricted to Podo- and Myovirus morphologies, respectively, while *N. multiformis* ATCC 25196 phages showed Myo-, Sipho-, and Podovirus morphologies. One lysate (LA) had an apparent filamentous phage; however, we could not assemble the genome for a filamentous or ssDNA phage from the obtained sequencing reads.

Infection passages were performed for each mixed phage lysate against each of the four AOB hosts for a preliminary assessment of host range. Infected cultures demonstrating early cessation or inhibition of nitrification were subjected to polymerase chain reaction (PCR) using phage-specific primers to confirm the presence of different phages capable of infection (Fig. 2). 42 different phage genomes were identified, with 16 of them showing potential infectivity against more than one host. Of these, 10 phages were capable of infecting hosts from both *Nitrosomonas* and *Nitrosospira*, five phages were capable of cross-genus infection but did not infect the other strain of the same genus, and one phage (rov1) could infect all four hosts (Fig. 2E).

**Figure 2.**
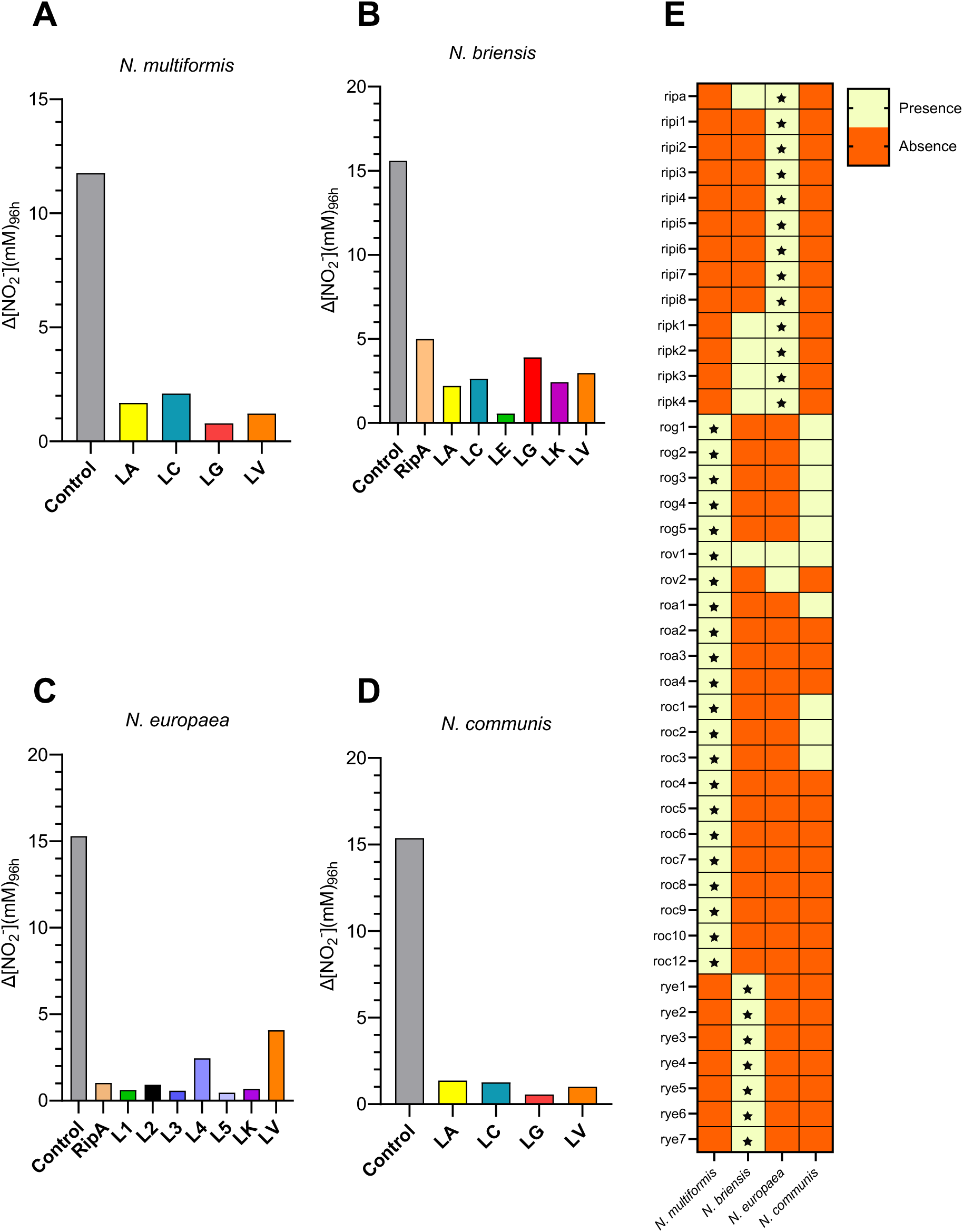
Host range of 12 cell lysates against four AOB species. A-D displays the difference in nitrite production after a 96-hour incubation, including an untreated control, depicted on the far left; (A) *Nitrosospira multiformis* ATCC 25196, (B) *Nitrosospira briensis* C-128, (C) *Nitrosomonas europaea* ATCC 19718, (D) *Nitrosomonas communis* Nm2. (E) Host range of specific phages within mixed lysates was confirmed by PCR using phage-specific primers. Absence of a band for a given phage, designated by an orange box, was deemed to result from no infection. Presence of a band for a given phage, marked with a wheat-coloured box, was deemeds to indicate successful phage amplification. Stars indicate phages matched to their isolation host.

### AOB phage genome features

Following Illumina sequencing of the 12 cell lysates, 45 dsDNA phage genomes were assembled and considered to be of high confidence based on assembly quality and viral hallmark gene content. Genomes ranged in size from the largest jumbo phage (roc1; 338.7 kB), to the smallest phage (ripi7_b; 12.7 kB) (Table S1). 11 of these phages were designated as putative temperate based on the presence of genes encoding integrases, with assembled genomes lacking annotated integrases marked as putative strictly lytic (Table S1). None of the temperate phage genomes were mapped back to the host genomes used in this study, indicating that they were obtained from the environmental screening and were not induced from the hosts during the isolation process. Temperate phage genomes were significantly different from their putative strictly lytic counterparts in %GC (p<0.0001) and genome size (p=0.0117) (Fig. S3). Most phage genomes showed no significant similarities to previously isolated or characterized phages, apart from a group sharing similarity to phage ΦNF-1, the only characterized phage of AOB which infects *Nitrosomonas europaea* ATCC 25978, *Nitrosomonas nitrosa* DSM 28438, and *Nitrosomonas communis* DSM 28436^12^. However, several temperate phages and proviral-like genomes were similar to proviral regions from other bacterial genomes (Table S3). Two closely related phages roc2 and ripk1 shared 97.35% similarity across >80% of their genomes to *Stenotrophomonas indicatrix* DR953 (CP118899.1), potentially representing a recent crossover event and an undescribed prophage region.

### Taxonomic diversity of AOB phages

The AOB phage genomes were classified using a multi-layered and hierarchical taxonomic approach. Clustering with GRAViTy v2 organized the genomes into 21 distinct ≥family-level clades (Fig. 3A). 13 phages grouped into orphaned monophyletic clades, with eight clades encompassing the remaining genomes. Clade 39 containing 10 genomes shared close homology (<0.4 distance threshold) to *Nitrosomonas* phage ΦNF-1 in the family *Autoscriptoviridae*. In our host range experiments, these phages only infected *N. europaea* ATCC 19718, while ΦNF-1 was shown to also infect *N. communis*^12^. Clade 31 was associated with the *Kyanoviridae*; however, it forms a deep branch and new family in *Pantevenvirales* when compared to *Kyanoviridae*, *Ackermannviridae* and *Straboviridae*, which are the three established families of *Pantevenirales*, and represents a sister family to *Kyanoviridae* (Fig. S4). Moreover proteins from these phages had <40% identity across all of the hallmark proteins of *Kyanoviridae* (Fig. S4)^17^. Clade 31 contained six distinct genera and tentatively given the family name *Araviridae*, order *Pantevenvirales* (Fig. 3B). Agreement between VipTree and vContact2 programs suggested Clade 31 contained two phages (rye2 and rog3) (Fig S5; Table S3). Orphaned monophyletic clades had insufficient representation to confidently assign higher-level taxonomy; thus, we propose six new viral families with some likely representing new orders. These are: *Araviridae*, *Druantiviridae*, *Cernuviridae*, *Sinaviridae*, *Fulfuniviridae*, and *Nantoviridae* (Table S4).

**Figure 3:**
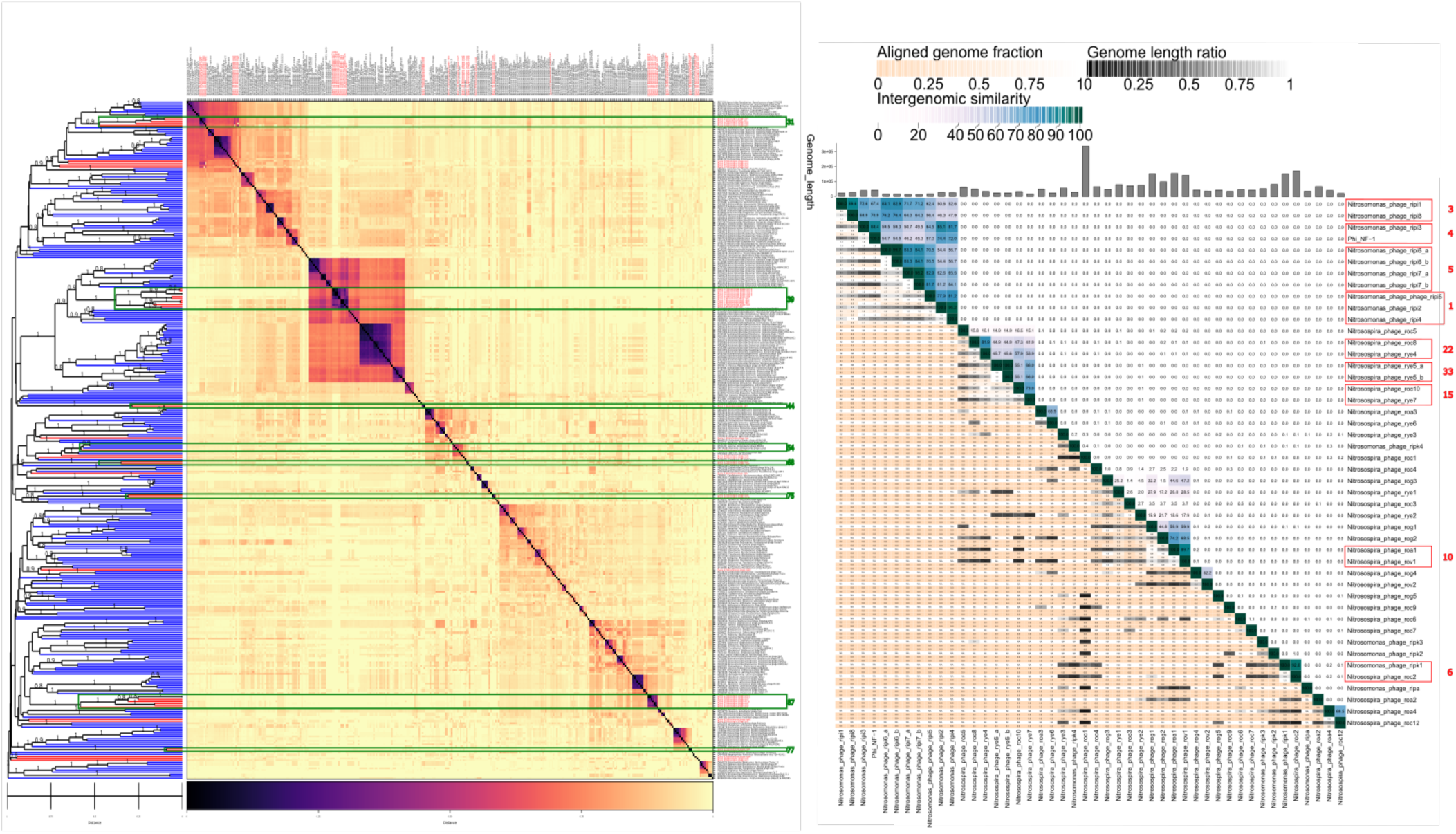
**(A)** GRAViTy v2 Family-level taxonomic distribution of 45 dsDNA AOB phage genomes with 300 reference ICTV genomes. Heatmap of proteome-wide similarity with darker colors representing greater similarity and lighter colors (wheat) representing low to no similarity. The dendrogram on the left shows reference genomes in blue, and query genomes in red, with leaf annotations on the right. Green boxes are numbered according to the assigned taxonomic cluster, marking clusters with multiple query representatives at the family-level. **(B)** Genus and species-level clustering of 45 dsDNA AOB phages using the VIRIDIC algorithm. The top bar graph shows the genome length with the intergenomic similarity marked with cool colors on the upper portion of the heatmap, and the lower portion represents the aligned genome fraction. Red boxes are labeled according to the number of their assigned genus-level clusters.

To further separate families into viral genera and species, VIRIDIC was used with 95% ANI species- and 70% ANI genus-level similarity cutoffs (Fig. 2B). In total, 34 genera were assigned, with nine containing ≥2 species. Phages belonging to the family *Autoscriptoviridae* represented four distinct genera, with *N. europaea* ATCC 19718 phage ripi3 being the only one belonging to *Catalonvirus*, while being a distinct species compared to ΦNF-1, with 88.4% intergenomic similarity. Although *Nitrosomonas* phages ripi1-8 share substantial similarity to ΦNF-1, they were 19.74 ± 7.89 kB smaller (Fig. S6). Moreover, the recent study characterizing ΦNF-1 demonstrated its successful infection of four *Nitrosomonas* hosts, including *N. communis* Nm2, which was also tested in this study^12^. However, no lysates containing *Autoscriptoviridae* phages could infect *N. communis* Nm2 in our experiments. Additionally, six *Nitrosomonas* phages (ripi6_a, ripi6_b, ripi7_a, ripi7_b, ripi2, ripi4) and two *Nitrosospira* phages (rye5_a, rye5_b) clustered as the same species (Fig. 3B; Table S4). A total consensus taxonomy was generated for all phage genomes as outlined in Table S4, with taxon names representing nature deities.

### Morons and auxiliary metabolic genes in AOB phage genomes

The ability of temperate phages to confer beneficial adaptations to their hosts has been extensively demonstrated in the literature^18–20^. However, the interaction between phages and nitrifying hosts has not been characterized outside of metagenomic inferences and limited data from ΦNF-1^12,21^. We anticipated that some putative temperate phages could have the capacity to modulate nitrification and alter host physiology. In this work, many phage genome-encoded auxiliary metabolic genes (AMGs) escaped initial sequence-based annotations, but were recovered using Phold, leveraging structural homology to annotate hypothetical proteins. Structural annotations were further confirmed through sequence-based methods. The Kyoto Encyclopedia of Genes and Genomes (KEGG) database was also used to match proteins to potential physiological and metabolic roles (Fig. 4E). A cumulative, but non-exhaustive, summary of auxiliary genes of interest found in this study is listed in Table S6.

**Figure 4:**
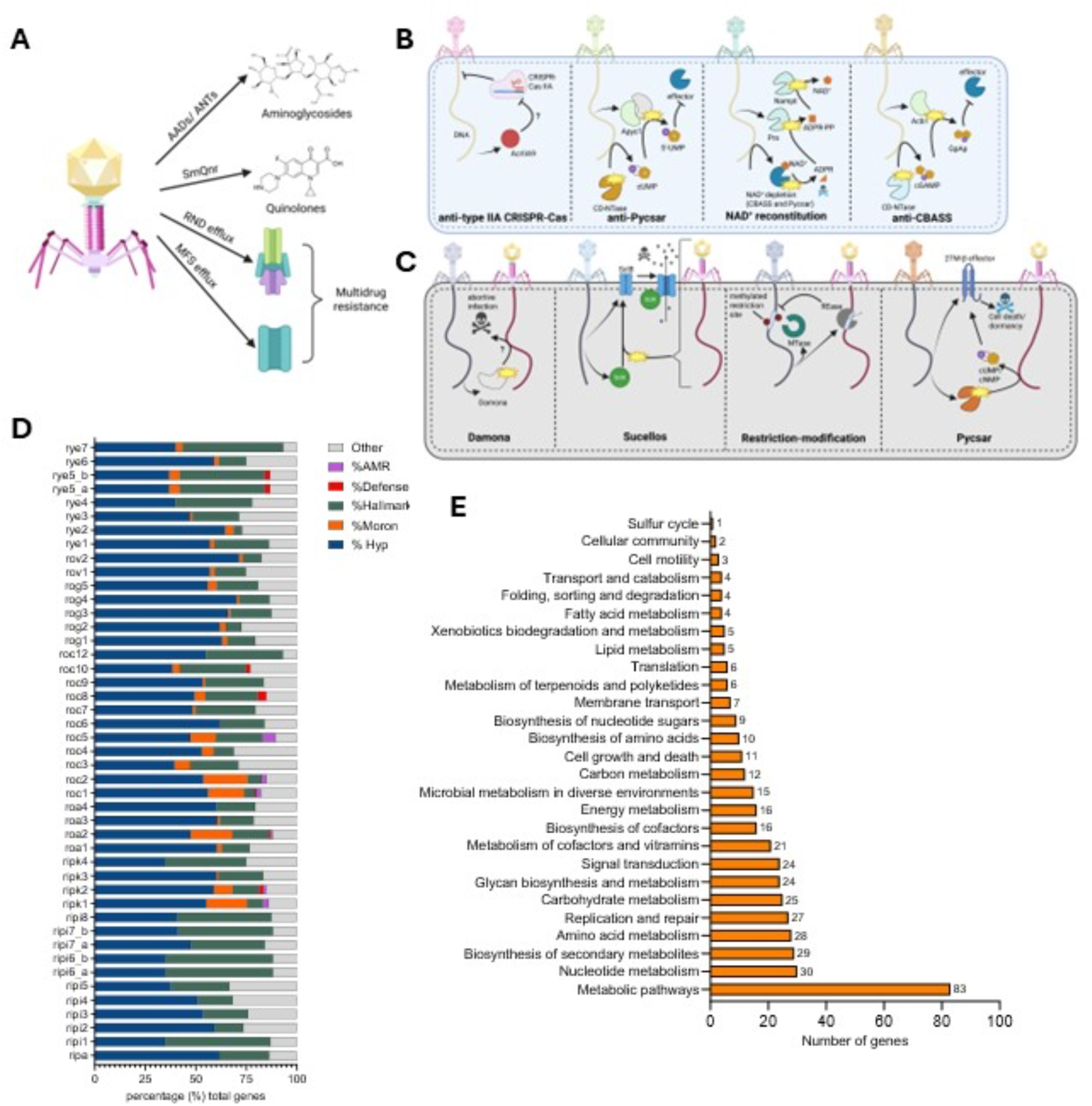
**(A)** Summary of antimicrobial genes encoded by phages infecting AOB: aminoglycoside 6-adeylyltransferases (AADs), aminoglycoside nucleotidyltransferase (ANT), resistance-nodulation-cell division (RND) efflux, and major-facilitator superfamily (MFS) efflux. **(B)** Counter-defense strategies encoded by AOB phages: type IIA anti-CRISPR-Cas (AcrIIA9) inhibition of defense/ recognition through what remains an unknown pathway^22^. Anti-Pycsar (Apyc1) cyclic pyrimidine monophosphate signaling molecule degradation and effector inhibition^23^. NAD^+^ reconstitution pathways (NARP1/2) to subvert abortive infection through NAD^+^ depletion as employed by Pycsar, CBASS, and other systems^24^. Anti-CBASS (Acb1) functions by linearizing cGAMP to disrupt signal reception by effector proteins^23^. **(C)** Anti-phage defense systems encoded by AOB phages: Damona remains poorly understood but induces an abortive infection response during viral infection^25^. Sucellose encodes a sensor SclA protein that activates SclB ion channels upon viral infection, resulting in membrane instability and abortive infection^25^. Type II restriction modification methyltransferases (MTases) methylate and mask restriction sites, preventing recognition and cleavage by restriction endonucleases (REases)^26^. Pycsar defense is encoded, producing cUMP/ cNMP signals that activate transmembrane (TM) effectors, resulting in growth arrest or cell death^23^. **(D)** Relative proportion of gene categories encoded in each AOB phage genome as percent (%) of total genes: antimicrobial resistance (AMR; purple), defense and anti-defense related proteins (red), hallmark proteins (green), auxiliary metabolic genes (morons; orange), and hypothetical proteins (Hyp; blue). **(E)** The total number of genes related to various metabolic pathways across all AOB phages as annotated by KEGG.

Proteins related to bacterial motility and signaling were identified in *Nitrosospira* phage roc1 encoding two flagellin A (FlaA) proteins, FlaG, FliJ, and a FliI ATPase (Table S6). Additionally, four *Nitrosospira* phages (roa1, rog1, rog2, roc3) encode for ambiguous flagellar proteins. *Nitrosospira* phage roc2 encoded the ATPase and PilT components of a PilT/PilU type IVa pilus. Additionally, phages roc1 and ripk2 encoded chemotaxis-related proteins, including CheR, CheY, and a methyl-accepting chemotaxis protein (Table S6). Two component system proteins, including response regulators, histidine kinases, and transcription factors, were well represented throughout the phage genomes (Table S6).

Transporter-associated proteins were amongst the most abundant of the auxiliary genes identified. 17 incomplete transporter systems were identified in different phage genomes, including ATP-binding domains, RND efflux domains, and phosphate transporter subunits (Table S6). Single-subunit transporters were widespread, with four phages (roc1, roc2, ripk1, and ripk2) encoding MFS transporters, and two *Nitrosospira* phages (roc10, rye7) encoding CorA family magnesium transporters. Two phage genomes (rip1 and roc2) encoded OmpA-OmpF family porins, acting in solute transport, phage receptors, and adhesion amongst many other properties^27,28^.

Interestingly, phage roc5 encoded a complete ABC transporter cassette with an arabinose-like solute-binding protein, two permeases (one carbohydrate), and an ATP-binding protein (Table S6). AlphaFold3 construction (pTM= 0.82, ipTM= 0.83) of the cassette, including membrane embedment, revealed a solute binding protein, dimeric ATP-binding domain, and a heterodimer permease domain for a 198.65 kDa structure (Fig S7). Moreover, an arabinose 5-phosphate isomerase was identified three genes upstream of the ABC transporter, suggesting that arabinose is a likely substrate (Table S6). To our knowledge, this represents the first phage genome reported to encode a complete ABC transporter, including metagenome/virome-assembled genomes (MAGs).

Many AMGs encoded in this dataset share a link to iron and other pertinent ions, both in transport and utilization. Notably, proteins such as hydroxylamine oxidoreductase (HAO) and cytochromes rely on ferrous hemes for function, serving a pivotal role in the hosts physiology^29,30^. Six TonB-dependent transporters were identified, which may also serve to import iron, heme, and other compounds that play pivotal roles in AOB physiology; TonB-dependent transporters may also serve a secondary role as phage receptors (Table S6)^31,32^. Notably, TonB-dependent siderophore receptor family proteins were encoded by ripk1 and roc2. Phages ripk2 and roc1 encoded for heme/hemoglobin family TonB-dependent receptors.

Temperate phage genomes also contained proteins likely involved in AOB metabolic modulation (Table S7). For example, the *Nitrosospira* roc1 genome encoded a substantial amount of such proteins, including a carbonic anhydrase (Table S6 & S7), and others having roles in iron utilization, including bacterioferritin for iron storage^33^, coproporphyrinogen III oxidase for heme biosynthesis^34^, a complete ubiquinol-cytochrome *c* reductase complex proteins (Fe-S subunit, b subunit, and c1 subunit), and NifU-like Fe-S cluster assembly protein (Table S6 & S7; Fig. S8). The roc1 genome also encodes cytochrome c-type biogenesis proteins CcmEFH (Table S6 & S7)

Additionally, proteins implicated in glycolysis were encoded by two *Nitrosospira* phages (roa2, roc1) including a phosphoglucomutase, class II fructose biphosphate aldolase, phosphoglycerate kinase, glyceraldehyde 3-phosphate dehydrogenase, and a pyruvate orthophosphate dikinase (Table S6 & S7). Enzymes for the pentose phosphate pathway, including the rate-limiting enzyme UDP glucose-6-dehydrogenase, were encoded in 7 phages (roc1, rog1, rog2, rov1, rye1, roa1, roa2) (Table S6 & S7). Together, these data suggest that many AOB phages, particularly those with a putative temperate lifecycle, encode a multitude of auxiliary genes implicated in host fitness, including key metabolic enzymes seldom found in other phage genomes. Moreover, there appears to be a clear link between iron/heme acquisition through dedicated transporters, and the presence of iron-dependent metabolic enzymes (Table S6). This is likely to play to the native hosts environment, with phages like roc1 encoded iron-dependent enzymes without a dedicated transporter may aid those in iron-replete environments, while those encoding siderophore receptors may facilitate improved fitness under iron-deplete conditions.

### Antimicrobial resistance, defense and antidefense systems in phage and host genomes

Wastewater influent serves as a particularly strong reservoir of antimicrobials and thus is a hotspot for antimicrobial resistance gene (ARG) mobilization^35^. Moreover, due to the abundance of prokaryotes and their cognate viruses in wastewater, we sought to understand the abundance of ARGs, and defense-related proteins encoded by AOB phages. The *N. briensis* C-128 and *N. multiformis* ATCC 25196 genomes both encode *adeF* for fluoroquinolone/ tetracycline efflux and *vanH* for glycopeptide resistance^36,37^ (Table S6). *N. europaea* ATCC 19718 and *N. communis* Nm2 similarly encode *adeF*, *vanT*, and *qacJ* for quaternary ammonium compounds in their genomes^37–39^. *Nitrosomonas* phage ripk1 carries a SmQnr family pentapeptide repeat protein implicated in quinolone resistance^40^, and ripk2 encodes a putative glyoxalase/ bleomycin resistance protein. Moreover, two resistance-nodulation-division (RND) efflux pumps (adaptor and permease) as well as four major facilitator superfamily (MFS) transporters are encoded in phages infecting both *Nitrosospira* and *Nitrosomonas* (Table S6). However, these transporters are known to have functions beyond antimicrobial resistance^41,42^ and thus can only be considered as conferring putative antimicrobial resistance. Although the functionality of these systems has not yet been demonstrated, these data suggest that phages may be playing a pivotal role in conferring newly acquired ARGs to their AOB hosts, providing an additional advantage in wastewater treatment and other environments exposed to antibiotics, including fertilized agricultural soils where AOB are particularly active^43^.

Due to the high degree of virus-host interactions in wastewater, bacteria and their cognate phages commonly interact through an arms race between defense and antidefense systems ^44^. Both *Nitrosomonas* and *Nitrosospira* host genomes encode extensive repertoires of defense systems that potentially subvert viral infection including type I-IV restriction modification (RM), type IC/F and III CRISPR-Cas, as well as type I and II cyclic oligonucleotide-based anti-phage signaling systems (CBASS), toxin-antitoxins, abortive infection (Abi) systems, and others (Table S8). Numerous defense-related proteins were identified in the phage genomes (Table S9), including the Damona and Sucellos defense systems which are encoded by the roc10 *Nitrosomonas* putative lytic phage and the ripk2 *Nitrosospira* temperate phage, respectively. Several antidefense systems were identified in the phage genomes, including NAD^+^ reconstitution pathway (NARP) proteins 1 and 2, which counteract NAD^+^ cleavage by the host to induce cell death. This pathway is initiated by numerous defenses including CBASS type II/III^24,45^. Additionally, *N. europaea* ATCC 19718 phage ripk1 encodes for an anti-CBASS 2 (Acb2) protein, known to be a cyclic-oligonucleotide signal receptor to counteract all known CBASS signals^46^. Moreover, ripk1 encodes for an anti-Pycsar protein (Apyc1) while the *N. europaea* ATCC 19718 genome lacks a Pycsar system. Of all publicly available *Nitrosospira* and *Nitrosomonas* genomes, *Nitrosomonas* sp. PRO4 (GCA_021405195.1) and *Nitrosospira* sp. time.spades.CONCOCT.1kb_041 encode Pycsar defense systems cognate to antidefense Pycsar systems encoded by ripk1 (Table S9). Pycsar defense systems are rare, with only these two instances amongst 3,058 defense and anti-defense systems encoded across available *Nitrosomonas* and *Nitrosospira* genomes. On the other hand, roc8 was found to encode for a Pycsar system itself, with a CBASS-like double transmembrane domain effector, and a Pycsar cyclase, which catalyzes the cyclic pyrimidine signaling needed for effector activation and abortive infection^47^. This suggests at least some of the AOB phages are likely to have broader host ranges than demonstrated here and could infect the cultivated AOB hosts used in this study even though they lack Pycar systems. There are also instances where defense and antidefense systems match between host and phage (Table S8; Table S9), indicating long-term evolutionary relationships between AOB hosts and cognate phages.

### Presence of phage genomes in metagenomic datasets

Prior to screening for AOB phages, it was anticipated that there would be a low abundance of phages infecting nitrifying hosts compared to other viruses, even in wastewater where AOB are enriched for nitrogen removal. Thus, to provide evidence on the abundance of these phages both in soil and wastewater environments, we screened publicly available short-read indiscriminate metaviromes from agricultural soil (478), and wastewater samples (676) (Table S10). Metavirome reads were mapped against assembled phage genomes to determine the presence of relatives (≥70% identity & ≥ 1x coverage) in the environment. Only *Nitrosomonas* phages ripk1, ripk2, and ripk4 and *Nitrosospira* phages roc1, roc2, and rog2 were found in soil metaviromes (Fig. 6). Phages ripk1, roc1, and roc2 were the most widespread phages across the samples screened, while ripk2, ripk4, and rog2 were seldom present, with relatives in 2, 1, and 2 metaviromes, respectively (Table S13; Fig. 4). Additionally, the most widespread relatives showed average breadths of 89.37% (ripk1), 81.63% (roc1), and 89.86% (roc2), being above or near genus-level identity across most samples. Moreover, total coverage was on average 14.04x ± 8.72.

**Figure 5.**
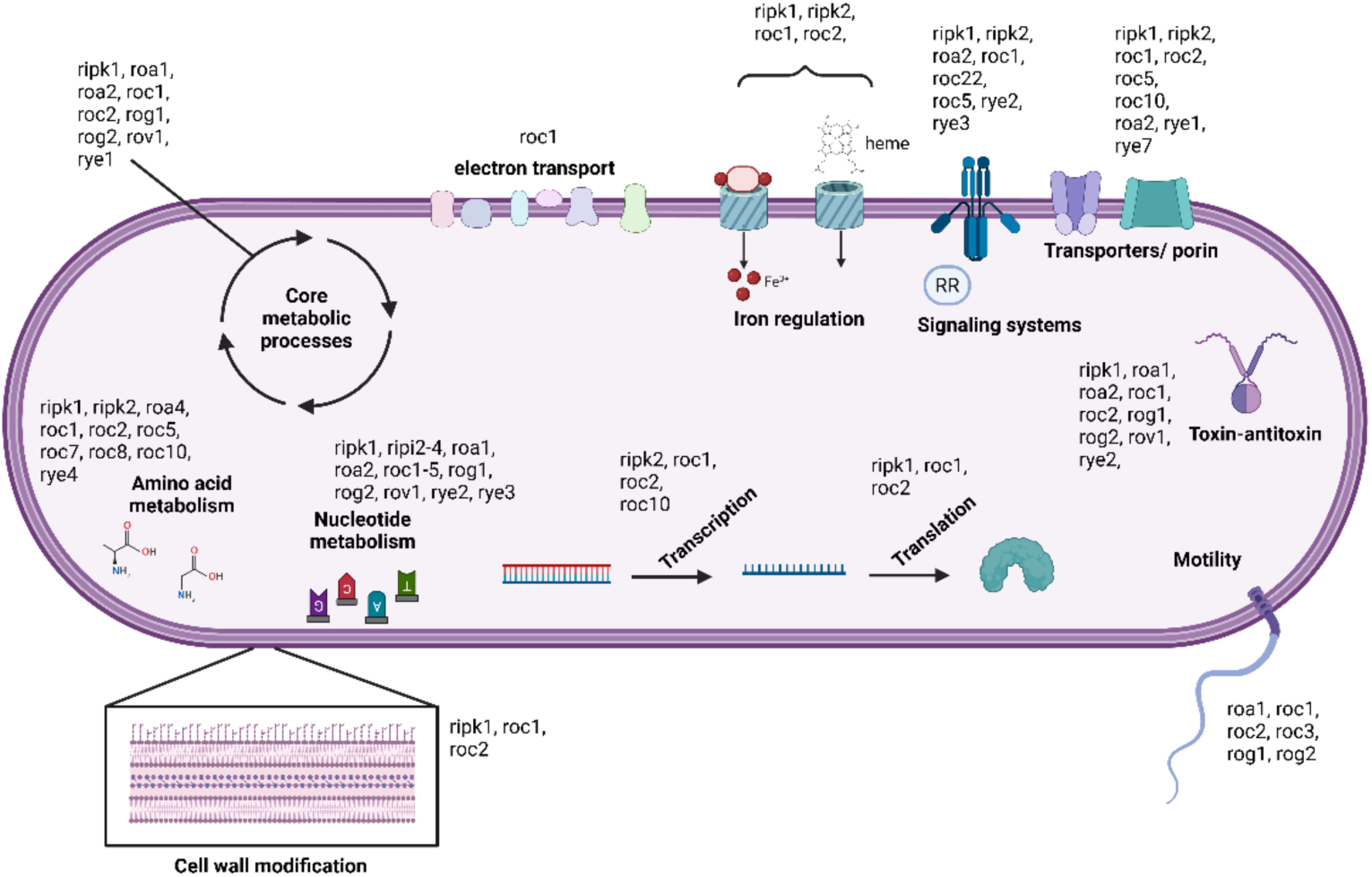
Non-exhaustive summary of auxiliary metabolic genes and morons encoded by AOB phages. Annotations were retrieved from KEGG, Phold and Pharokka. Bolded text indicates the functional category, with phages encoding related proteins in un-bolded text. Specific proteins of interest can be found in supplementary tables S6 and S7.

**Figure 6:**
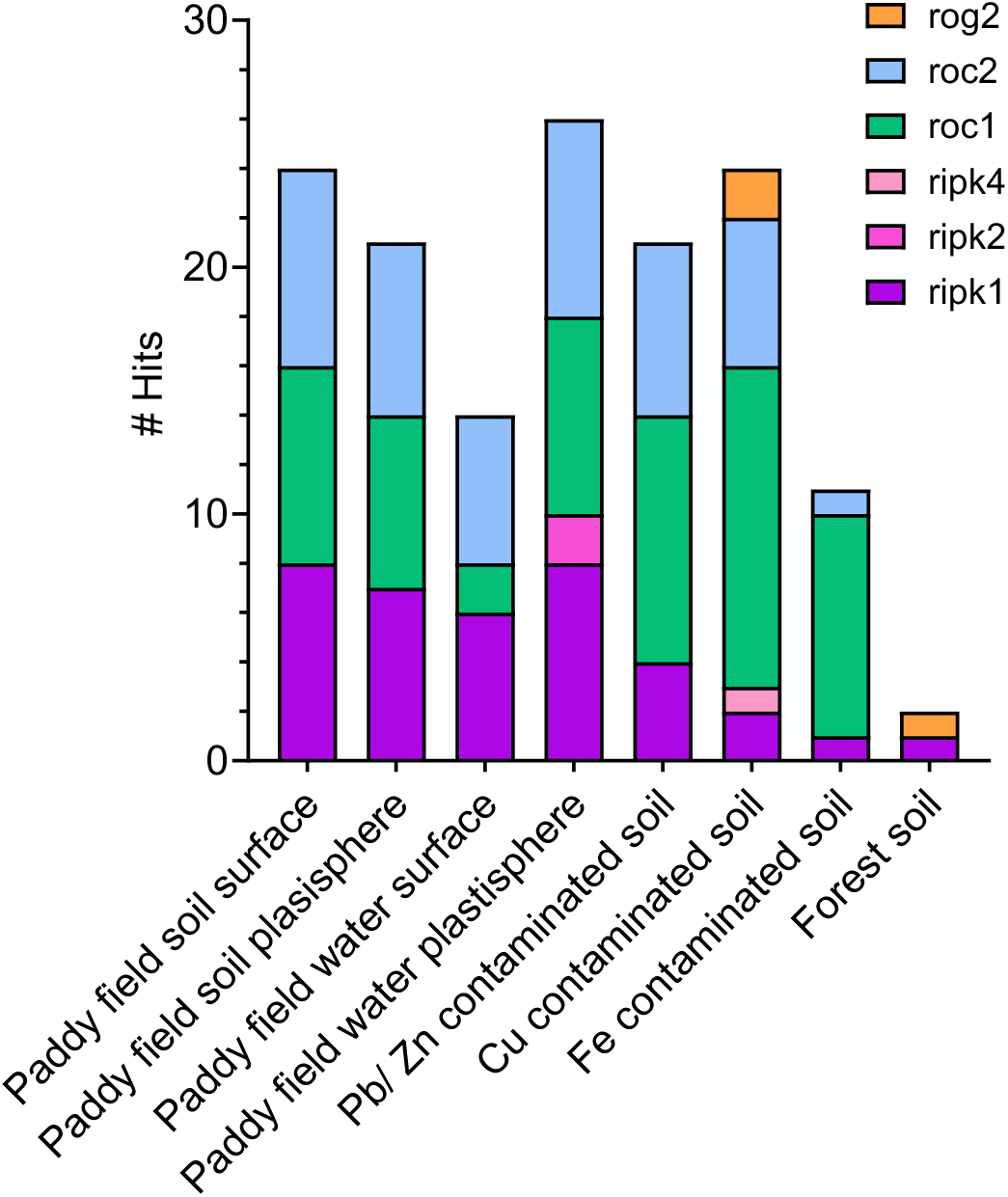
Number of hits for AOB phages identified in this study in soil metaviromes. Illumina-sequenced metaviromes from soil-associated samples were screened for ≥1x coverage and ≥70% breadth of the total query genomes. Hits were categorized based on sample site conditions and environment, with the total number of metavirome hits for each phage retrieved on the Y-axis.

Surprisingly, when surveying the 676 wastewater metaviromes from 63 cities across the globe (PRJEB87273^45^), no substantial coverage across the majority of AOB phage genomes from this work was identified. We anticipate that this may have been a result of AOB phages being extremely low in relative abundance compared to other viruses in a rich and diverse environment as wastewater treatment. These data suggest that the recovery of AOB phages has not only avoided extensive culturing and retrieval, but their diversity has likely also been neglected in metagenomics. Due to the limited hits recovered, relative-abundance metrics could not be calculated; however, the RPK and coverage statistics are shown in Figure S9.

## DISCUSSION

Fundamental knowledge of AOB phages has been limited up to now due to the low population density of AOB and their cognate phages in natural systems and the incapacity, so far, to grow AOB on solid media, precluding normal plaquing assays used in the isolation and characterization of novel phages. Using a filtration technique and liquid cultivation method, we identified and characterized 45 dsDNA phages and their genomes. Not only could this method recover numerous phages in mixed lysates but can produce pure lysates of target phages (*Nitrosomonas* phages ripa and ripi1-4). AOB phages were diverse, encompassing the three main *Caudoviricetes* phage morphologies and clustering into six novel families and 41 genera. Phage-containing lysates inhibited nitrification in liquid AOB cultures. 11 putative temperate and 20 putative strictly lytic phages carried AMGs, and morons implicated in altering host physiology, and most phage genomes encoded ARGs and/ or defense/antidefense systems. The significant early cessation of nitrite production observed from mixed AOB phage provides a strong basis for the eventual use of cocktails of AOB phages to inhibit nitrification in ecosystems suffering from nitrogen oversaturation and N_2_O emissions, such as agricultural soils. In many biocontrol and phage therapy applications, cocktails of at least three phages have been shown to minimize host immunity^48^, which is likely true for AOB. The use of a broad-host-range phage is most likely to achieve a more complete impact. However, the deployment of temperate phages that encode ancillary genes implicated in elevating host fitness are not desirable for deployment in open ecosystems as this could increase nitrification activity over time. Moreover, overlap between target community phage defense and phage-encoded anti-defense systems, which would provide a distinct advantage to phage cocktails, should be considered. Combining phage cocktails in combination with fertilizers would enable AOB growth, which is necessary to enable phage infection, and provide an effective method of phage dissemination. However, assessing viral stability in liquid fertilizer formulations, optimizing phage stabilizers for long-term survival, assessing the efficacy of phage in controlling the activity of AOB in biofilms, and pilot-scale proof-of-concept deployment are necessary before large-scale interventions.

## MATERIALS AND METHODS

### Growth of ammonia-oxidizing bacteria

Three strains of AOB: *N. multiformis* ATCC 25196, *N. briensis* C-128, and *N. communis* Nm2 were grown in HEPES-buffered HK medium at pH 7.7 and 10 mM (NH_4_)_2_SO_4_ with 0.5% (v/v) Phenol red as a pH indicator^38^. *Nitrosomonas europaea* ATCC 19718 cultures were grown in ATCC medium 2265. Cultures were grown at 28°C on a 120-rpm orbital shaker (New Brunswick Scientific) in 125- or 250-mL Wheaton bottles with inlayed butyl rubber stopper caps. pH was maintained at ∼7.5-8.0 using 5% (w/v) Na_2_CO_3_. Growth was measured via nitrite production using a colorimetric nitrite assay until cultures converted ∼90% of the ammonia substrate to nitrite^49^. A 2-4% (v/v) inoculum of near-stationary culture was used to transfer cultures; however, for phage propagation, a 20% (v/v) stationary inoculum was used with 2-4% (v/v) inoculum of 0.45 µm-syringe filtered (Biosharp, mixed cellulose ester) phage lysate.

### Wastewater samples preparation

Raw influent samples were collected from two urban wastewater treatment plants in Alberta, Canada. Filtrates were prepared by passing samples over 0.45-µm pore size cellulose acetate filters (Sartorius) to remove contaminants and produce the filtered influent samples. All filtration steps were completed using a Büchner-style vacuum filtration setup with a 1000-mL collection flask, a sand core glass membrane base, and a 500-mL filter cup.

### Vacuum filtration screening for phages infecting AOB

To screen for phages infecting AOB, a modified filtration method from Ghugare and colleagues (2017) was used, where ca. 1-5×10^9^ cells were captured on 0.45-µm pore size cellulose acetate filters (Sartorius) as determined by direct cell counts using a hemocytometer and phase-contrast light microscopy (Motic B5 professional series). 100-250 mL of filtered influent was passed through filters on which host cells were previously deposited, and the filtrate was recovered for re-use on cell-deposited filters containing other host cells. Host cell-deposited filters were transferred to 25 mL of AOB cultures in growth medium, and nitrite production was measured over 7 days (as described above), as a time course series from the same sample for each phage/ lysate. Cultures demonstrating a cessation in nitrite production and significantly lower nitrite yield compared to untreated control cultures were transferred ≥5 times before further phage analysis to eliminate the potential for non-infectious viral carryover. Control cultures underwent the same steps with sterile AOB medium replacing filtered wastewater samples.

Infection-positive cultures were filtered once more through 0.45-µm pore size syringe filters to remove unlysed cells. To concentrate phage lysates, 15-mL 100 kDa Amicon tubes were used following the phage-on-tap protocol (Bonilla et al., 2016). Lysates were 100-fold concentrated in SM buffer and transferred to 1.5 mL microcentrifuge tubes following the phage on tap concentration and wash ultrafiltration protocol using 100 kDa Amicon Ultra-15 tubes (Millipore)^50^. All concentrated lysates were stored at 4°C until further manipulations. Published methods for AOB phage isolation including the addition of raw influent filtrate to mid-log AOB cultures, and the addition of polyethylene glycol 8000 (BioShop) to raw influent filtrate, were attempted, but neither method resulted in detectable reduction of nitrite production. The filtration technique further enabled the passage of the same raw influent filtrate to screen against all AOB hosts in sequence.

### Transmission electron microscopy

5 µL of 100x concentrated lysates in SM buffer were applied to 300 mesh Ted Pella formvar/carbon coated copper grids (Sigma-Aldrich), allowed to sit for 1 minute, and stained with 10 µL 4% uranyl acetate (Fisher Scientific) for ∼10 seconds. Images were taken using a Phillips/ FEI (Morgagni) transmission electron microscope with a Gatan camera.

### Phage validation by PCR

Host range analysis was performed by propagation of phages against the four AOB hosts. The propagation of phages targeting AOB hosts was conducted over ≥5 passages before samples were recovered for PCR using primers specific to each phage (Supplementary Data Sheet 1). To achieve PCR amplification of low-abundance phage fragments, Q5 hot start polymerase was used following the manufacturer’s instructions with a 65°C annealing temperature for 35 cycles. Bands were visualized in 1% (w/v) agarose gels with 1x Tris-acetate EDTA buffer at 100 V for ∼50 minutes.

### DNA extraction and genome assembly

DNA was extracted from concentrated phage lysates using the Qiagen Viral RNA kit. Samples were submitted to and sequenced by Genome Québec using Illumina NovaSeq paried-end 150 bp 35M reads. Read quality was assessed using FastQC (Andrews, 2010). Before assembly, reads were mapped to host genomes (*Nitrosospira multiformis* ATCC25196 (GCF_000196355.1), *Nitrosospira briensis* C-128 (GCF_000619905.2)*, Nitrosomonas europaea* ATCC19718 (GCF_000009145.1) using BWA-MEM2 v2.3 to remove host contamination^51^. Genomes were assembled thereafter using SPAdes v4.2.0^52^ (--meta, --only-assembler). Contigs underwent two rounds of polishing using Pilon v1.24^53^. The origin of phage contigs was assessed using a combination of CheckV^54^ (reference data 1.5, Galaxy version 1.0.3+galaxy0), VIBRANT^55^ (Galaxy version 1.2.1+galaxy2), and Virsorter2^56^(database v0.4, Galaxy version 2.2.4+galaxy0) using default parameters on the Galaxy EU webserver^53^, selecting candidate contigs that met a ≥ medium quality threshold, or contained ≥3 hallmark phage genes. To further confirm that reads were not host-contaminated, <u>BLASTn</u> was used to compare viral contigs against host genomes and plasmids.

### Genome annotation

Candidate contigs were annotated using Pharokka^57^ v1.9.0 using default parameters with –dnaapler for genome reorientation based on the terminase large subunit (TerL) when possible. A list of proteins identified as hypothetical, that may have escaped annotation was curated through a structure-guided approach using Phold^58^ v1.2.0 (GPU, default parameters). As Phold assigns ambiguous names to annotated proteins (e.g., VFDB virulence factor), hits were searched using the virulence factor database (VFDB) v6.0^59^, DefenseFinder^60–62^ v2.0.1, BlastKOALA^63^ and AcrFinder^64^ for more precise annotations, and BLASTp was performed (ClusteredNR database) for all previously hypothetical proteins with an annotation cutoff of 40% identity and 80% query coverage with an e-value < 1e-5. Protein re-annotations are recorded in supplementary Table S6. To determine the domains of particularly meaningful proteins discussed in the main text, Interproscan^65^ (https://www.ebi.ac.uk/interpro/) and PFamScan^66^ (https://www.ebi.ac.uk/jdispatcher/pfa/pfamscan) were leveraged using default parameters.

### Phage taxonomy

To evaluate the taxonomy of AOB phages and propose novel taxa, a hierarchical approach was used to generate consensus assignments and groupings of viral genomes. Family-level classification and infer potential order-level branches, GRAViTy v2^67^ version 2.2 and ViPTree^68^ v4.0 were used with support by vContact2^69^ v0.12.0 (default settings) to determine proteome-based clustering of phages with known ICTV *Caudoviricetes* phages. GRAViTy v2 results took precedence over other tools; however, VipTree and vContact2 were also used to reveal consensus and disagreement clusters. VipTree phylogenetic trees were refined in iTOL (https://itol.embl.de/). GRAViTy v2 was used with 300 reference *Caudoviricetes* genomes from the 2024 ICTV release^70^. Taxon assignment was done using the “full classification new” pipeline with default parameters in VipTree and vContact2. Consensus family-level assignment was determined based on the agreement of 2/3 programs. To refine the assignment, genus- and species-level clustering was performed using VIRIDIC^71^ v1.1 with 95% and 70% similarity thresholds for species and genus, respectively. Additionally, Clinker^72^ (40% identity cutoff, default settings) was used on the CAGECAT web server^73^ to visualize genomic similarities between genus members, highlighting conserved and variable regions.

### Metavirome screening

To determine the presence of AOB phages in both soil and wastewater metaviromes, a total of 478 soil-associated and 676 wastewater viromes were searched using BBMap v39.52^74^ with ≥ 70% identity and average fold ≥ 1x cutoffs following adaptor trimming with fastp v1.0.1^75^. Reads per kilobase (RPK) was computed by the average read coverage divided by the contig length in kb. All unfiltered hits against soil and wastewater metaviromes, along with their SRR accessions, are available in Tables S10 and S11, respectively.

### Statistical methods

Statistical analysis and graph preparation of experimental data were performed using GraphPad Prism 10. No data were excluded from analyses. All code was run using Python 3.11.9 and Conda v 25.9.1.

## Data availability

Genome assemblies for all phage genomes in this study have been deposited in GenBank and will be available upon publication under the Bioproject PRJNA1457874. Individual genome accessions can be found in Supplementary Table S1.

## Code availability

No custom code was developed for this study. All analyses were performed using Visual Studio Code on Windows Subsystem for Linux 2 (Ubuntu) for programs outlined in the methods, with any non-default parameters outlined in the methods.

## ACKNOWLEDGEMENTS

LYS and DS were funded by the Natural Sciences and Engineering Council of Canada Discovery Grants Program. LYS was funded by the Canada Research Chairs program and the Grantham Foundation. AM was funded by Medical Research Council **CLIMB-BIG-DATA (**MR/T030062/1).

## AUTHOR INFORMATION

### Authors and Affiliations

**Department of Biological Sciences, University of Alberta, Edmonton, AB, Canada**

Lisa Y. Stein and Aaron A.B. Turner

**Department of Genetics and Genome Biology, University of Leicester, United Kingdom**

Andrew Millard

**Department of Chemical and Materials Engineering, University of Alberta, Edmonton, AB, Canada**

Dominic Sauvageau and Miranda Stahn

### Contributions

M.S performed filtration optimization and recovery efficiency. A.A.B.T recovered the phage lysates and performed all other experiments as well as sequencing and data analysis. A.A.B.T conceived and designed the experiments under the guidance of L.Y.S, D.S, and A.M. A.A.B.T, L.Y.S, D.S, and A.M wrote the manuscript.

## ADDITIONAL INFORMATION

Supplementary Information is available for this paper

Peer review information includes the names of reviewers who agree to be cited and is completed by Nature staff during proofing.

Reprints and permissions information is available at www.nature.com/reprints

## Notes

### Competing Interest Statement

The authors have declared no competing interest.

### Summary of Updates

The title of the article has been changed to better reflect the findings

