## Supplementary material for "Diversity of Novel Bacteriophages Infecting Ammonia-Oxidizing Bacteria": Supp. methods and figures

**Supplementary Methods:**

**Auxillary metabolic gene cataloging.** To retrieve a list of hypothetical proteins that could be annotated but escaped initial annotation by Pharokka, Phold was used for structure-guided annotation. However, many hits from Phold were ambiguous and assigned names based on database hits (e.g., VFDB virulence factor). To eliminate false hits and probe for more meaningful genetic nomenclature, sequence- and HMM-based tools were used on these hits, assigning names for those showing >40% protein identity and >80% coverage from tools like BLASTp. Additionally, sequence-based versions of the databases leveraged by Phold were used for more meaningful insight: virulence factor database (VFDB), CARD, DefenseFinder, and acrFinder. The subsequent putative proteins were catalogued with cognate locus tags.

**Transmission electron microscopy.** TEM was performed (Phillips/ Morgagni) with a Gatan camera. 5 uL of concentrated phage stock was applied to a gold grid using 1% (w/v) uranyl acetate. Images were taken at 110,000X magnification and analyzed using ImageJ (Schneider et al., 2012).

**Protein structure models.** To determine the metabolic role of encoded genes and potential implications for nitrification, KO assignment and annotation was performed using the "Blast KEGG Orthology And Links Annotation" (KOALA) webserver (https://www.kegg.jp/blastkoala/; Kanehisa et al., 2015). Moreover, the novelty of proteins was determined using the functionally and evolutionarily significant novel (FesNov) catalogue using EggNOG-mapper v2 (http://eggnog-mapper.embl.de/; Cantalapiedra et al., 2021; Rodríguez del Río et al., 2023). Proteins of particular interest were visualized using the AlphaFold3 web server (https://alphafoldserver.com/) with 20 unique iterations. To simulate the inner cell membrane to generate a more accurate structure of transporter complexes using PPM 3.0 (Lomize et al., 2021), structures were compared against known complexes using the FoldSeek-multimer search tool against all databases.

**Evaluation of filtration recovery.** To optimize the filtration protocol, phage T4 (ATCC 11303-T4) was used as a proxy to determine the recovery efficiency of low-abundance viruses. *Escherichia coli* ATCC 11303 was grown in Luria-Bertani (LB) medium to establish growth on solid medium and for liquid cultures. Cultures were grown at 37°C at 120 rpm following inoculation with 1% (v/v) mid-log culture. Growth was measured by optical density (OD) at 600 nm using a spectrophotometer. For T4 infection, the virus was added with a multiplicity of infection (MOI) of 0.01 once cell cultures reached 0.1-0.3 $OD_{600}$. Lysates were harvested using a 0.22-µm syringe filter. Titer was determined using a double-layer agar overlay (Kropinski et al., 2009) following a serial dilution. Untreated *E. coli* was used as a control, with undiluted T4 lysates used as a positive indication of infection (titers ranging from $10^9$-$10^{10}$ PFU/mL).

**Supplementary Figures:**

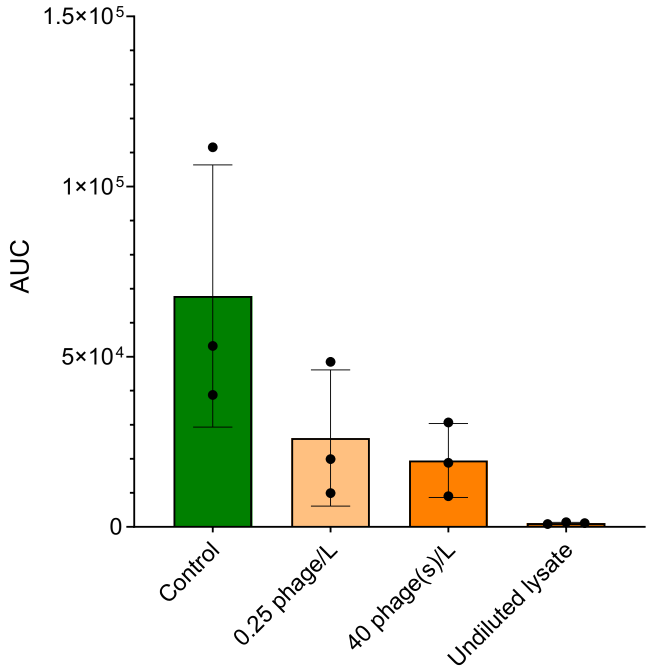


**Figure S1: Validation and sensitivity of cell-deposited filter based phage screening determined with *E. coli*-phage T4 system.** Area under the curve (AUC) of kill curves determined by $OD_{600}$ measurement following filtration-based screening of phage T4 infecting *E. coli* across 3 distinct replicates (n=3). Controls were untreated filters of *E. coli*. Treatments ranged in dilution: 0.25 phages/L (1 phage per 250mL sample), 40 phages/ L and undiluted lysates. Error bars represent the standard deviation in AUC values calculated across 3 replicates with individual values marked by black dots.

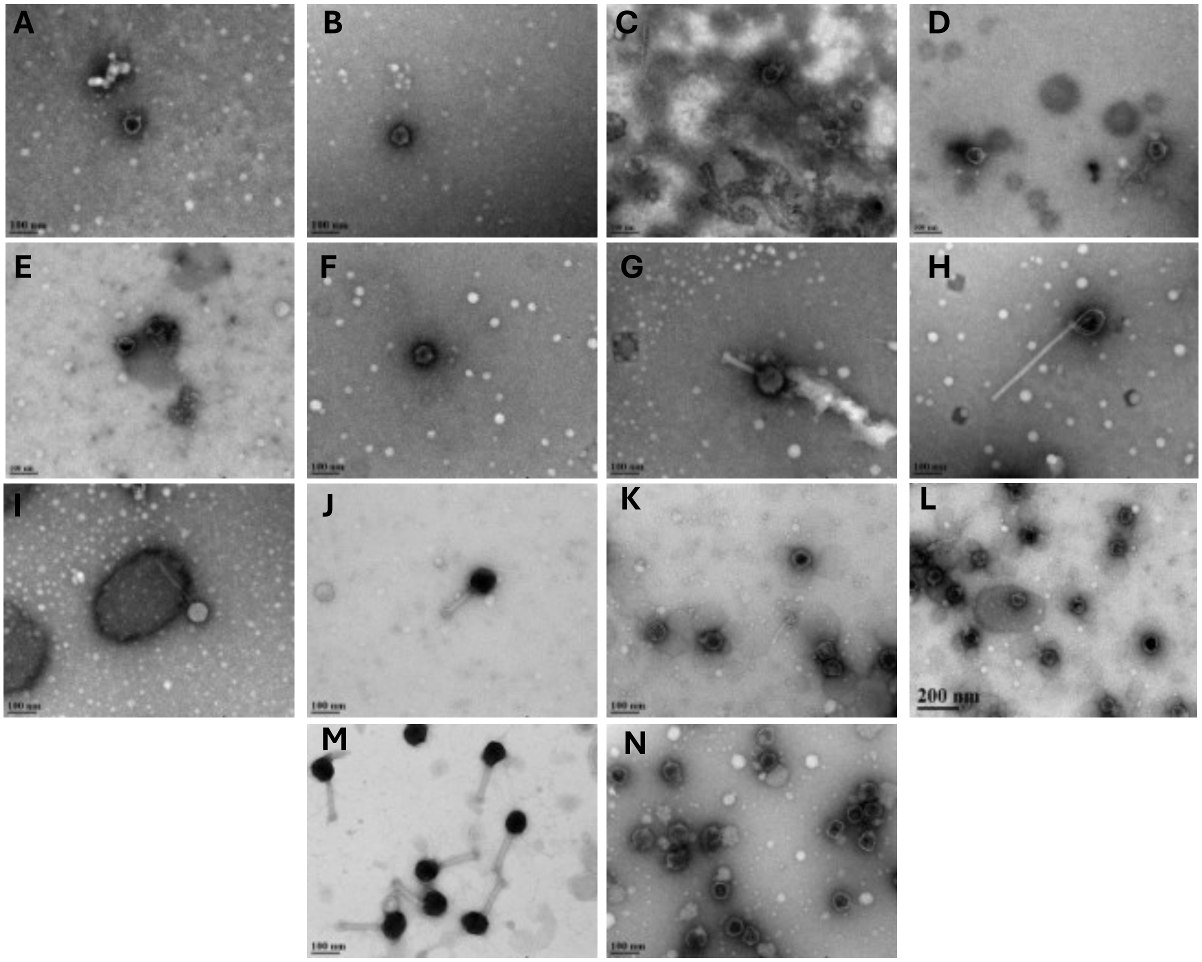


**Figure S2: Transmission electron microscopy of lysates collected** following sequential passaging of 12 lysates infecting AOB. *Nitrosomonas europaea* ATCC 19718 lyaste (**A**) RipA (*Nitrosomonas* phage ripa), (**B**) L1 (*Nitrosomonas* phage ripi1), (**C**) L4 (*Nitrosomonas* phage ripi4), (**D**) L3 (*Nitrosomonas* phage ripi3), (**E**) L2 (*Nitrosomonas* phage ripi2), (**F**) L5. *Nitrosospira multiformis* ATCC 25196 lysates (**G**) LG, (**H**) LG, (**I**) LG, (**J**) LA, (**K**) LC, and (**L**) LC, depicting a jumbo-like phage on the left. *Nitrosospira briensis* C-128 lysate (**M**) LE, and *N. europaea* lysate **(N)** LK.

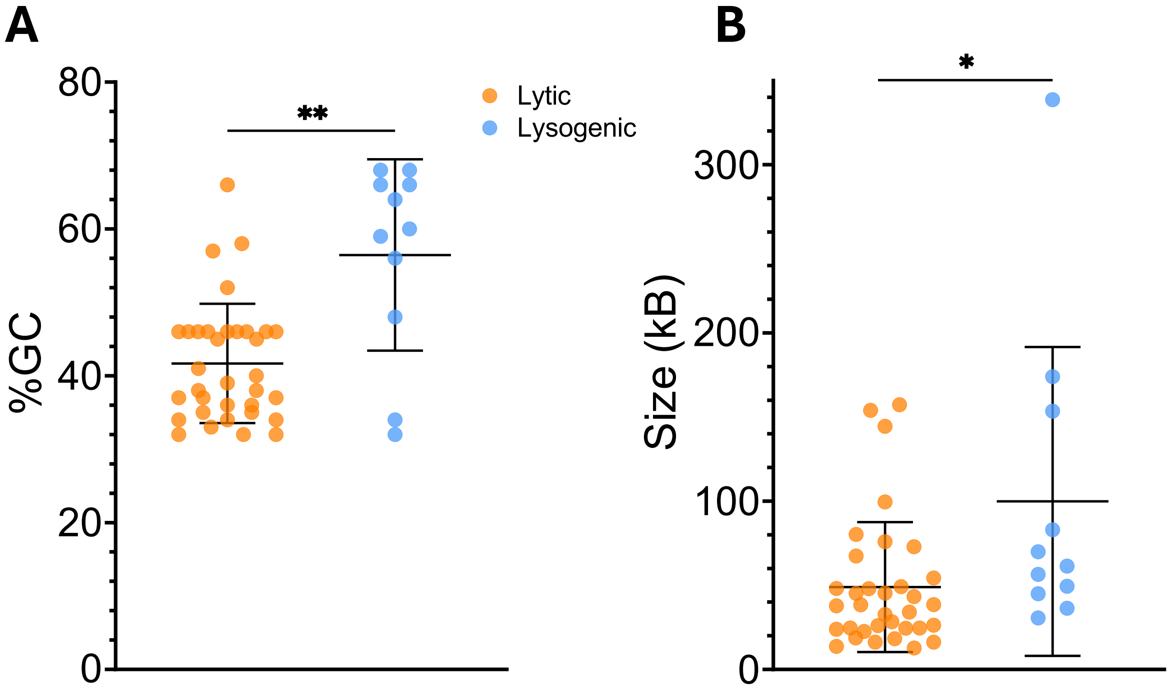


**Fig S3: Comparative genomics between putative strictly lytic (orange, n=34) and lysogenic (blue, n=11) AOB phages**: (**A**) percent GC content and (**B**) genome size. Statistical significance was determined using a two-tailed T-test and p-value is indicated by: p≤0.05 (*) or p≤0.01 (**).

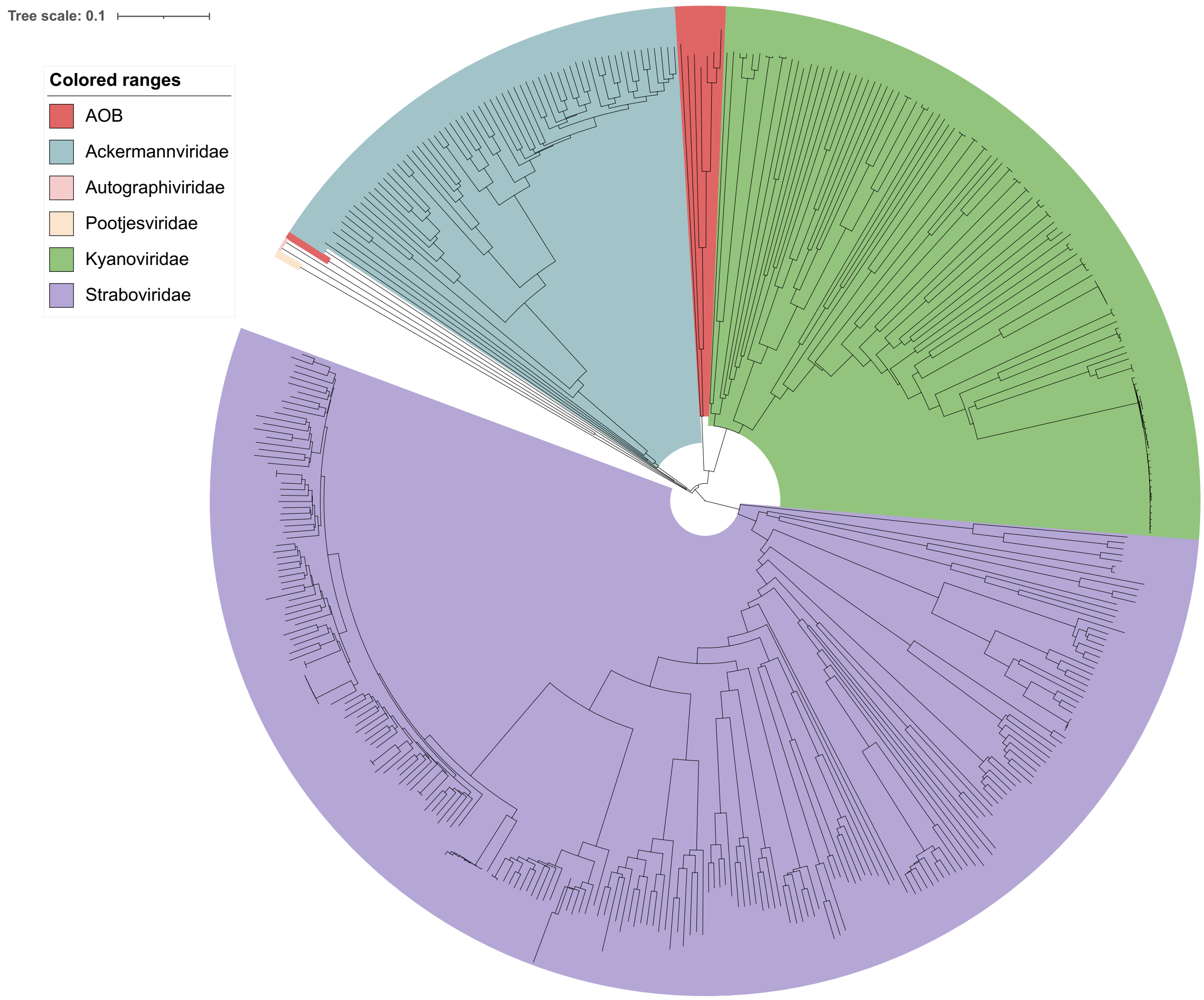


**Figure S4: Whole-proteome phylogenetic tree generated of *Pantevenvirales* AOB phages**. VipTree was used to compare *Pantevenvirales* AOB phages (red: roa1, rog1, rog2, rog3, rye1, rye2, rov1) to *Ackermannviridae* (blue), *Straboviridae* (purple), and *Kyanoviridae* (green). AOB phage rog4 was used as an outgroup along with representative phages from *Pootjesviridae* (yellow) and *Autographiviridae* (pink).

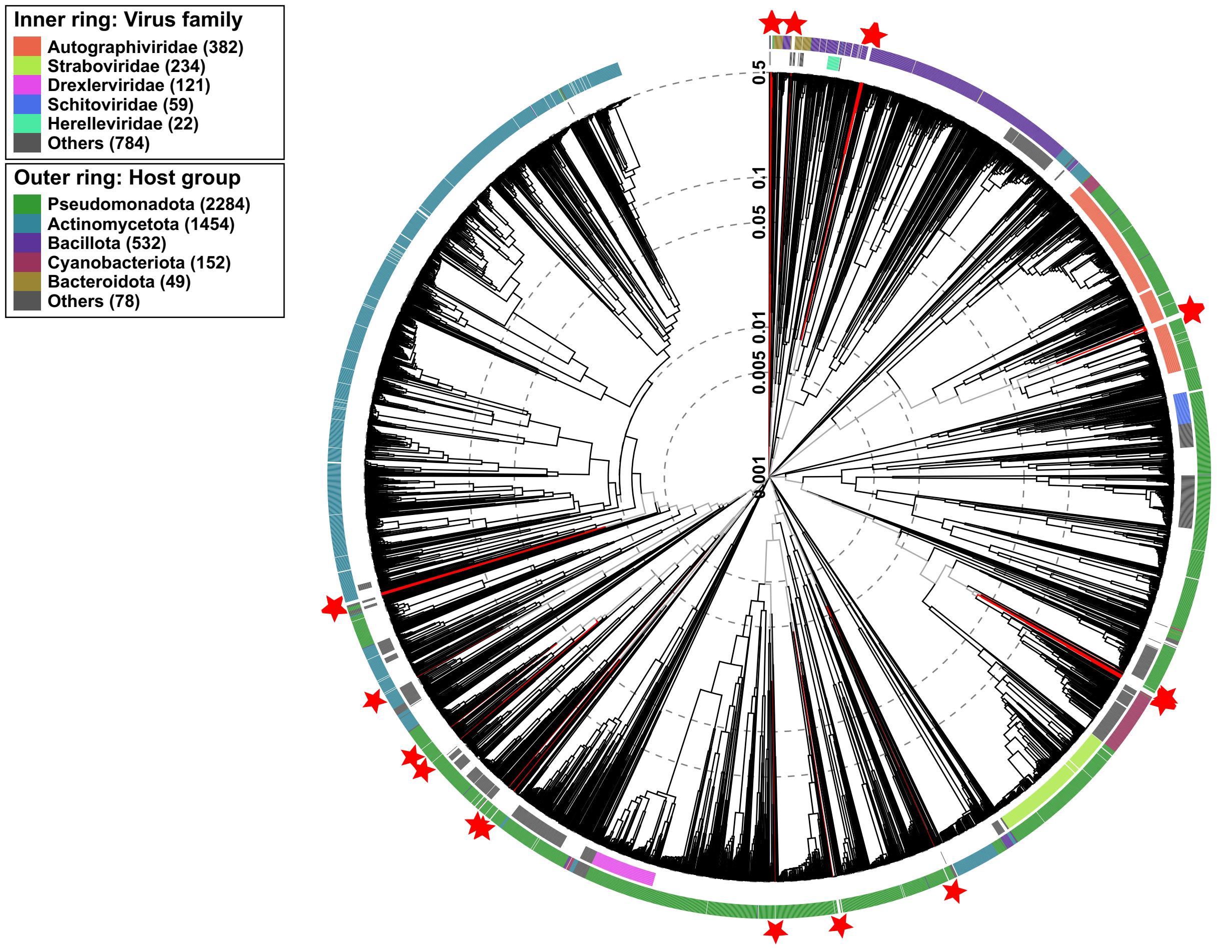


**Figure S5: Whole-proteome phylogeny of all AOB phages to known phages in the ICTV database using VipTree.** Inner rings designate known viral families, outer rings mark the infecting host. AOB phages are marked with red stars. Family-level clades are marked by a 0.05 log-scaled branch distance.

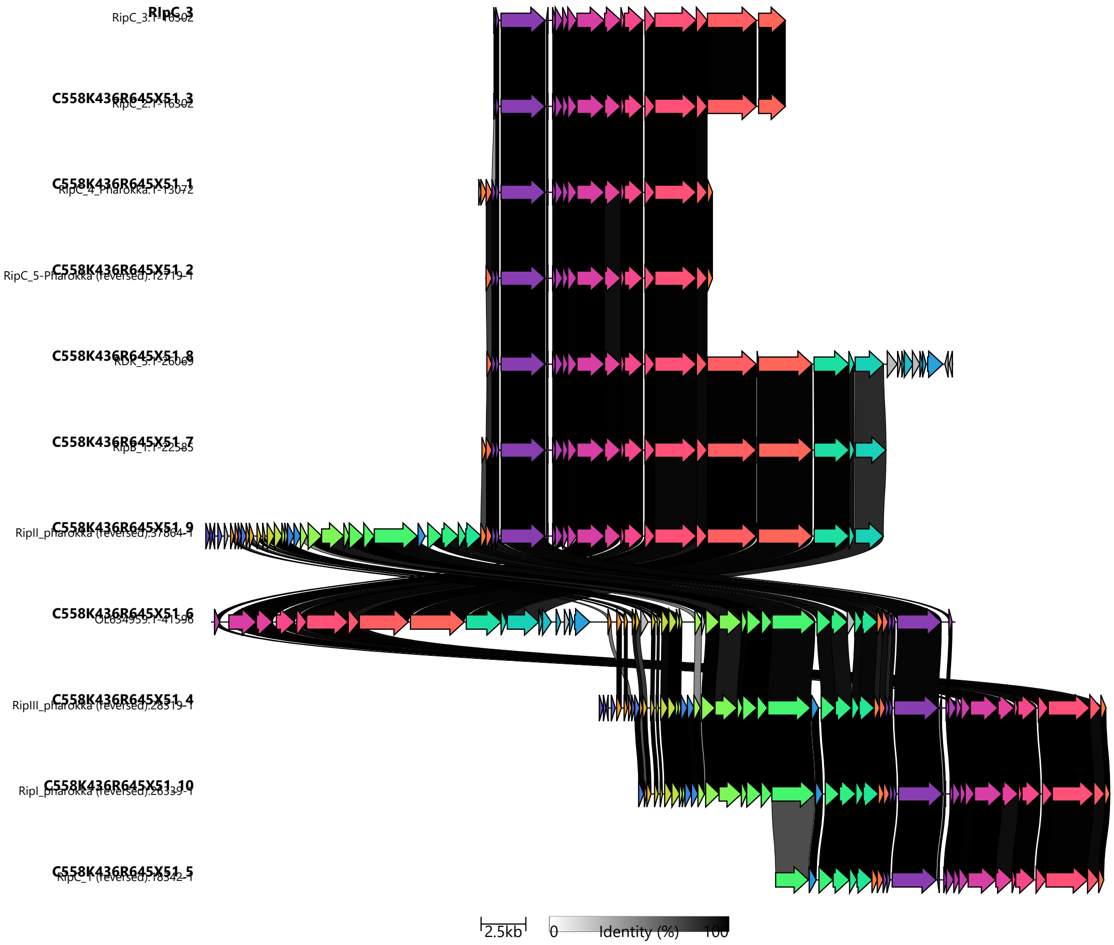


**Figure S6: Protein-sharing map of all proteins that shared ≥30% similarity of AOB phages belonging to *Autoscriptoviridae***, along with *Nitrosomonas* phage ΦNF-1.

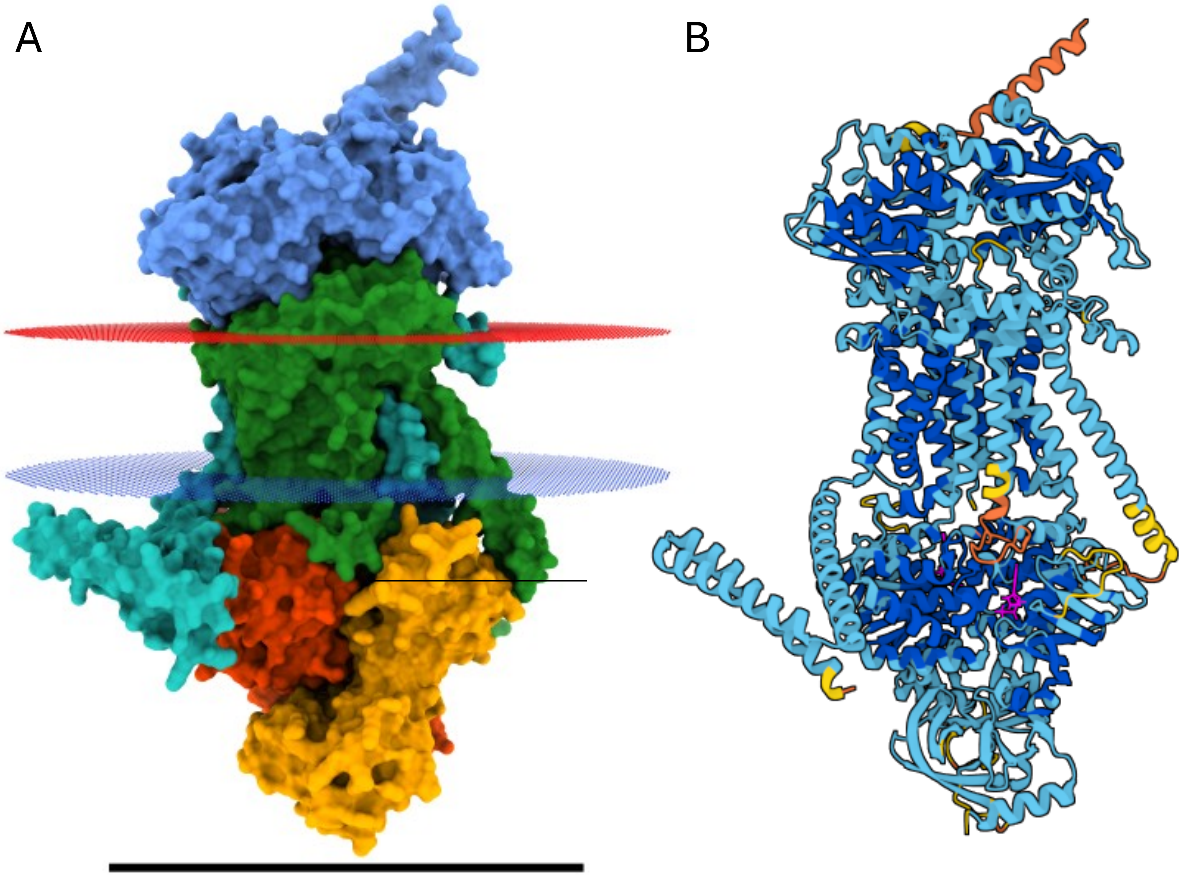


**Figure S7: AlphaFold3 construction of the ABC transporter complex from *Nitrosospira* phage roc5.** (**A**) Domain diagram depicting the arabinose-like domain solute-binding protein in blue, carbohydrate permease (teal), generic permease (green) and two copies of the ATP-binding domain (orange and red). Structure simulation in the inner membrane shows the periplasmic side in red and cytoplasmic in blue. A scalebar below shows a 100 Å distance. (**B**) A pLDDT map of the complex generated by AlphaFold3 with ATP bound (magenta) with a ipTM and pTM of 0.82. Structures were generated following 20 unique instances on the AlphaFold3 webserver, selecting the highest confidence structure generated (https://alphafoldserver.com).

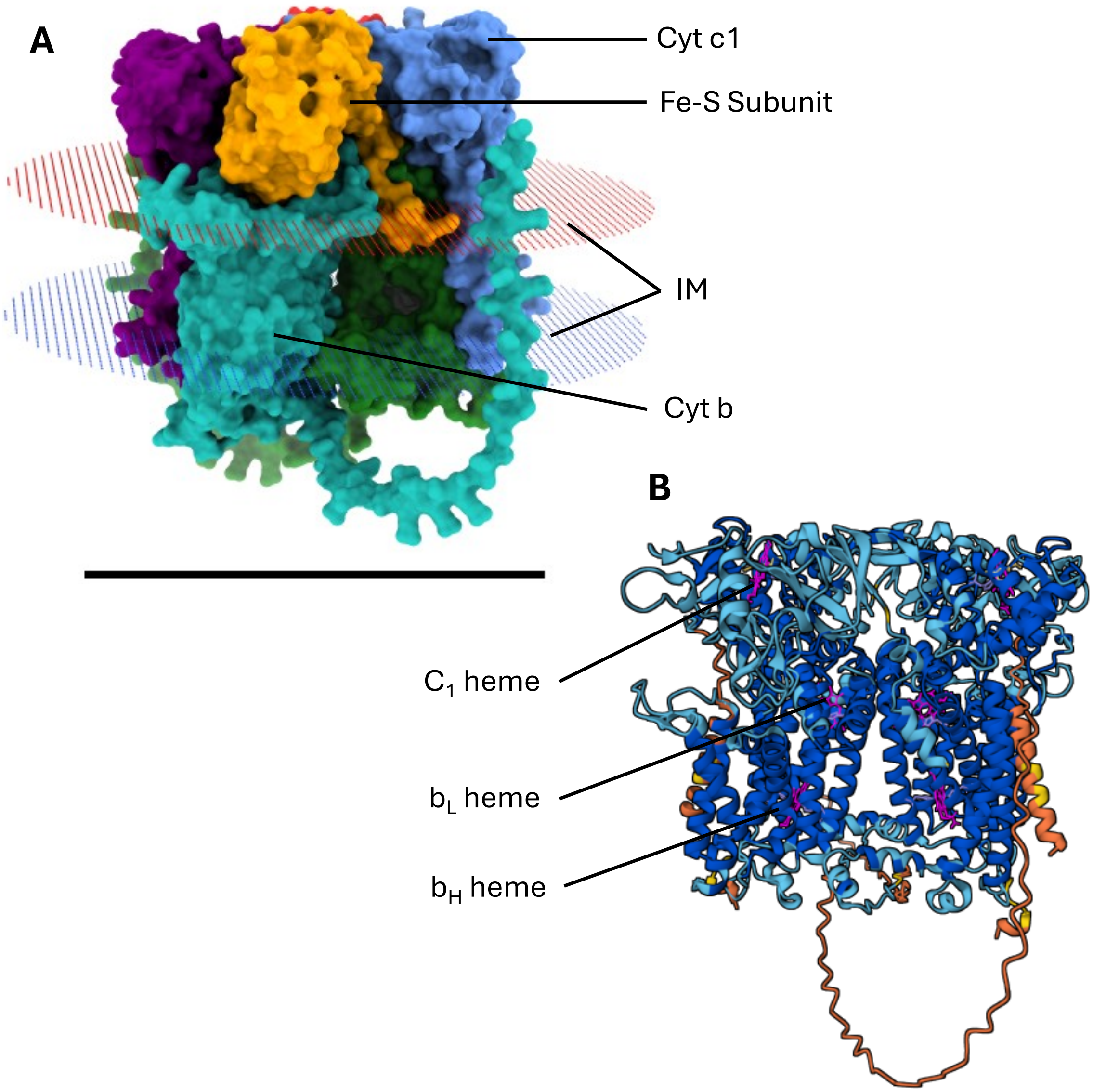


**Figure S8: AlphaFold3 construction of the ubiquinol cytochrome c reductase complex from *Nitrosospira* phage roc1.** (**A**) Domain diagram depicting the domains of the dimeric complex: cytochrome c1 subunit (blue and purple), the Fe-S subunit (orange and red), the cytochome b subunit (teal and green) in addition to the cytoplasmic (blue) and periplasmic (red) sides of the inner membrane (IM). A scalebar below shows a 100 Å distance. (**B**) A pLDDT map of the complex generated by AlphaFold3 with c and b-type hemes bound (magenta) with a ipTM and pTM of 0.87. Structures were generated following 20 unique instances on the AlphaFold3 webserver, selecting the highest confidence structure generated (https://alphafoldserver.com).

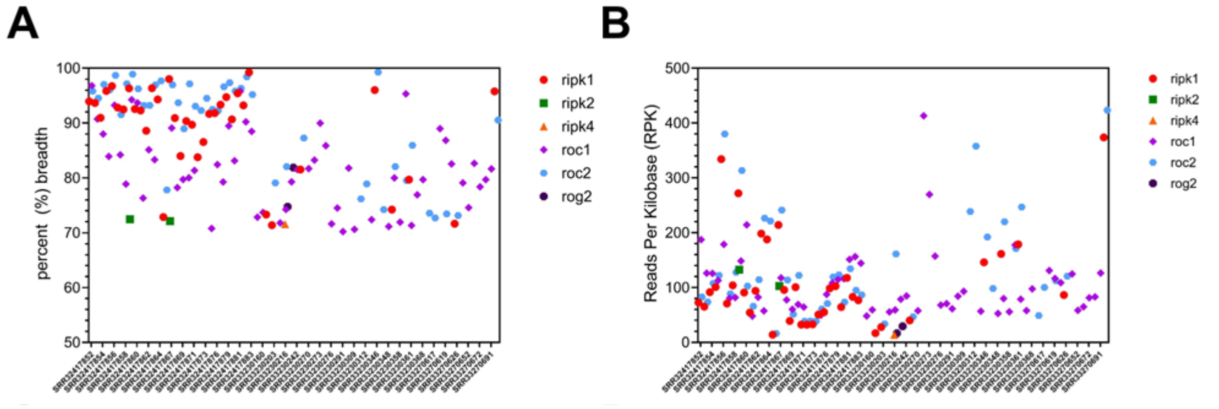


**Figure S9:** Metagenome analysis of all bacteriophages infecting ammonia-oxidizing bacteria. Illumina sequenced metagenomes from soil samples were screened for ≥1x coverage and ≥70% breadth of the total query genomes. (**A**) Total breadth percentage is shown for high-confidence mapping for soil with cognate metavirome accessions in the x-axis. (**B**) The relative abundance calculated by reads per kilobase (RPK) is shown on the y-axis.
